# Effects of imidacloprid and thiamethoxam at LC_30_ and LC_50_ on the life table of soybean aphid *Aphis glycines* (Hemiptera: Aphididae)

**DOI:** 10.1101/2021.04.06.438575

**Authors:** Aonan Zhang, Lin Zhu, Zhenghao Shi, Kuijun Zhao, Tianying Liu, Lanlan Han

**Author notes:** Corresponding author, (HL).

## Abstract

The soybean aphid, *Aphis glycines* Matsumura (Hemiptera: Aphididae), is a main pests of soybean that poses a serious threat to its production. Studies were conducted to understand effects of the different concentrations of the insecticides (imidacloprid and thiamethoxam) on the life table of *A. glycines* to provide vital information for its effective management. We found that the mean generation time, adult and total pre-oviposition periods in *A. glycines* specimens exposed to LC_50_ imidacloprid and thiamethoxam were significantly longer than those in the control group. However, when exposed to LC_30_ imidacloprid and thiamethoxam, the adult pre-ovipositional period was significantly shorter than that in the control group. The mean fecundity per female adult, net reproductive rate, intrinsic rate of increase, and finite rate of increase were significantly decreased in individuals exposed to LC_30_ and LC_50_ concentrations of imidacloprid and thiamethoxam, respectively (*P* < 0.05). Both insecticides produce stress effects on *A. glycines*, and specimens treated with LC_50_ concentrations of the two insecticides exhibited a significant decrease in their growth rates than those treated with LC_30_ concentrations. This study provides data that can be used as a reference to predict the effect of imidacloprid and thiamethoxam on the population dynamics in the field, and agricultural producers could attach importance to prevent stimulation the reproduction made by low-lethal concentrations during actually applying pesticides.

## Introduction

The soybean aphid, *Aphis glycines* Matsumura (Hemiptera: Aphididae), was first detected in North America in 2000 and had rapidly spread to north-central and northeastern United States and southeastern Canada. Although the damage from *A. glycines* to soybeans is rarely devastating in Asia, it is considered a primary pest in North America (Ragsdale, Voegtlin, & O’neil, 2004; Wang, Kritzman, Hershman, & Ghabrial, 2006; Ragsdale, Landis, Brodeur, Brodeur, & Desneux, 2011; Hopper et al., 2017).

Foliar insecticides have been commonly used to effectively control *A. glycines* (Lee, 2000). By the late 1990s, neonicotinoid insecticides had been introduced worldwide due to their high efficiency, low toxicity, and wide application range (Srigiriraju, Semtner, & Bloomquist, 2010; Basit, Saeed, Saleem, Denholm, & Shah, 2013). The sales of neonicotinoids accounted for more than 25 % of the global insecticide sales in 2014 (Bass, Denholm, Williamson, & Nauen, 2015). Imidacloprid is a first-generation neonicotinoid (Matsuda, Buckingham, Kleier, Rauh, Grauso, & Sattelle, 2001), and thiamethoxam is a second-generation neonicotinoid and has higher activity, better safety, and longer duration of efficacy (Shi, Wang, Liu, Qi, & Yu, 2017).

The insecticidal effect of a particular insecticide is closely related to its type, application method, and application frequency (Xie et al., 2010). The drift and degradation of neonicotinoids themselves in the fields would make the doses of some aphids exposed to lower then those normally recommended for control. The lower doses permit survival of heterozygotes for resistance thereby fixing a resistant gene in a population more rapidly (Nauen & Denholm, 2005). The risk of the rapid development of resistant populations, secondary pest outbreaks, and rapid deterioration of the ecological environment was greatly increased (Isman, 2006; Khan, Abbas, Shad, & Afzal, 2014; Hanson et al., 2017; Zhao et al., 2018; Somar, Zamani, & Alizadeh, 2019).

A monitoring study performed in 2008 and 2011 reported an increased adaptation of the cotton aphid *Aphis gossypii* Glover to neonicotinoids in the south-central United States (Gore et al., 2013). In addition, cotton aphids from six different locations in Korea also exhibited adaptation to common neonicotinoids (Koo, An, Park, Kim, & Kim, 2014). Imidacloprid was the first commercial neonicotinoid used to control rice planthoppers in the early 1990s; however, after 10 years of its large-scale use in Asia, the sensitivity of rice planthoppers to imidacloprid decreased (Matsumura et al., 2013; Tao et al., 2019). *Bemisia tabaci* biotype Q (Hemiptera: Aleyrodidae) showed a similar response to conventional neonicotinoids (Luo, Jones, Devine, Zhang, Denholm, Denholm, & Gorman, 2010; Nauen et al., 2014).

To solve the recent problem of the rapid increase in pest adaptation to neonicotinoids, it is of great significance to carry out population dynamic analysis and to formulate an accurate application scheme (Feliciangeli & Rabinovich, 1985; Stark & Wennergren, 1995; Gabre, Adham, & Chi, 2005; Bass et al., 2011; Abbas, Shad, & Shah, 2015; Jan, Abbas, Shad, & Saleem, 2015). In this study, based on the life table of *A. glycines*, we evaluated the effects of imidacloprid and thiamethoxam on the reproductive development and population dynamics of *A. glycines* to provide a reference for the accurate use of these insecticides for controlling *A. glycines* populations.

## Materials and methods

### Laboratory aphid population and chemical agents

The laboratory strain of *A. glycines* used in this study was originally collected from a soybean field in Harbin, Heilongjiang Province, China. This strain had been cultured in the laboratory for several years and never been exposed to any insecticides. Dongnong 52 soybean plants were used to maintain the strain of *A. glycines* at Northeast Agricultural University, China. Soybean plants were grown in pots (15 cm diameter × 17 cm depth), with six plants in each pot kept at 25 ± 1 °C, a relative humidity of 65–70%, and a photoperiod of 14:10 (L:D) h. The laboratory aphid colony was maintained in the same environmental conditions as the chamber used for plant germination. Twelve pots with soybean plants were placed in a large tray (70 × 60 cm; L × W). Twice a week, one third of the old aphid-infested soybean plants (i.e., the four oldest pots with an aphid infestation) were removed and replaced with new aphid-free ones. Aphids were transferred by placing infested leaves on uninfested plants. This prevented the accumulation of excessive honeydew and sooty mold and ensured the provision of a homogeneous soybean plant for the aphids to feed on (Menger et al., 2020).

Water dispersible granules of insecticides (70% imidacloprid and 50% thiamethoxam) were purchased from North China Pharmaceutical Group Corporation, Hebei, China and Shaanxi Thompson Biotechnology Co., Ltd., Shaanxi, China respectively. Calcium nitrate, potassium nitrate, potassium dihydrogen phosphate, magnesium sulfate, disodium ethylenediaminetetraacetic acid (disodium EDTA), and streptomycin sulfate were all purchased from Shanghai Alighting Biochemical Technology Co., Ltd., Shanghai, China.

### Preparation of culture medium

Non-toxic, transparent plastic Petri dishes (6 cm diameter × 1.5 cm height) were used to perform the bioassay on newly hatched nymphs of *A. glycines* and the life table study. The components of the plant nutrient solution concentrate used to prepare the medium were as follows: calcium nitrate (4.1 g), potassium nitrate (2.5 g), potassium dihydrogen phosphate (0.7 g), magnesium sulfate (0.6 g), 1.54% disodium EDTA aqueous solution (5.0 mL), one million units of streptomycin sulfate (0.05 g), and distilled water (5.0 L). The diluent was obtained by mixing the plant nutrient solution concentrate with distilled water at a ratio of 1:3. Agar was prepared by mixing 1% w/w agar powder with diluent and was boiled while constantly mixing. After cooling for approximately 10 minutes, the warm agar was poured into the Petri dishes to a depth of at least 3–4 mm. At least 10 mm distance was allowed between the top of the agar and the rim of the Petri dishes. The metal tube was used to cut leaf discs from clean, untreated leaves. The leaf discs were 2 mm lesser in diameter than the Petri dishes and were attached to the agar medium with the top-side facing down. A metal tube was sharpened and cleaned regularly to ensure the clean cutting of the leaf discs. *A. glycines* on leaf discs fed on the bottom surface. Each Petri dish was then placed upside down to keep *A. glycines* in a natural feeding state. The incision was kept neat to avoid excessive crushing of the tissue at the edge of the leaves when they were cut. This prevented the leaves from rapidly developing mildew.

### Dose response bioassay

The dose response bioassays were conducted with newly hatched *A. glycines* nymphs using a leaf dip method recommended by the Insecticide Resistance Action Committee (IRAC; http://www.irac-online.org/resources/methods.asp). Insecticidal stock solutions were prepared in 1% acetone and further diluted to different concentrations using distilled water containing 0.05 % (v/v) Triton X-100 before using in dose response bioassay. According to the preliminary bioassays, seven concentrations of imidacloprid (19.95 mg a.i./L, 13.70 mg a.i./L, 9.10 mg a.i./L, 6.10 mg a.i./L, 3.47 mg a.i./L, 2.35 mg a.i./L, 1.88 mg a.i./L) and thiamethoxam (29.95 mg a.i./L, 24.98 mg a.i./L, 14.97 mg a.i./L, 10.05 mg a.i./L, 4.94 mg a.i./L, 3.64 mg a.i./L, 1.98 mg a.i./L) were prepared respectively. Fresh soybean leaf discs were immersed in solutions of seven concentrations; each leaf disc was immersed in a specific concentration for 10 s, removed from the solution, and placed on paper towels (abaxial surface facing up) to air dry. The control leaf disc was immersed in a solution of distilled water containing 0.05 % (v/v) Triton X-100 and 1 % acetone. The air-dried leaf discs were attached to the agar medium with the top-side facing down and newly hatched nymphs were placed on them. Treatment details (insecticide, concentration, and date) were recorded for each Petri dish. A small drop of distilled water was placed on the surface of the agar prior to laying the leaf on the surface to help the leaf stick to the agar surface. Sixty newly hatched nymphs were used for dose response bioassays at each concentration; three replicates were used per concentration and each replicate had 20 newly hatched nymphs. Mortality was determined after 24 h of exposure. The newly hatched nymphs were considered dead if they were found upside down and not moving or if they did not move when prodded with a small paint brush (Cordero, Bloomquist, & Kuhar, 2007). The toxicity of imidacloprid and thiamethoxam to the nymphs were statistically analyzed using SPSS (version 23.0, SPSS Inc., Chicago, IL, USA) and the LC_50_ and LC_30_ values for the newly hatched nymphs were obtained.

### Life table study

One hundred and fifty apterous adults were transferred onto fifteen leaf discs using a small paint brush and 10 apterous adults were placed on each leaf disc. Each Petri dish containing a leaf disc was sealed with a close-fitting, ventilated lid. Newly hatched nymphs were selected 24 h later and placed on a leaf disc pre-impregnated with LC_30_ and LC_50_ imidacloprid and thiamethoxam or a leaf disc pre-impregnated with distilled water containing 0.05 % (v/v) Triton X-100 and 1 % acetone.

One hundred newly hatched nymphs were exposed to each treatment; each newly hatched nymph was kept in a separate Petri dish, and each Petri dish was treated as one replicate. The growth, survival, mortality, and fecundity of the individuals were observed until all organisms died. After reaching the adult stage, the number of newly hatched nymphs reproduced by each adult every day was recorded and removed after recording.

### Life table analysis

The age-stage-specific survival rate (*s_xj_, x* = age, *j* = stage), age-specific survival rate (*l_x_*), age-stage specific fecundity (*f_xj_*), and age-specific fecundity (*m_x_*) were calculated as follows (Chi & Liu, 1985; Chi & Getz, 1988; Chi, 1988).

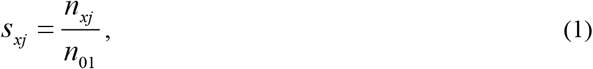

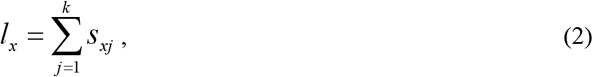

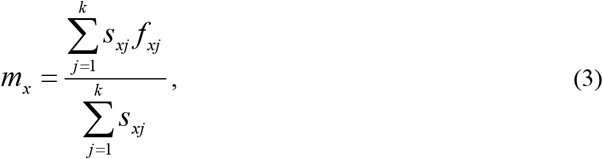

where *n*_01_ stands for the number of newly hatched nymphs and *k* stands for the number of stages. The net reproductive rate (*R*_0_), intrinsic rate of increase (*r*), finite rate of increase (*λ*), and mean generation time (*T*) were calculated as follows (Goodman, 1982):

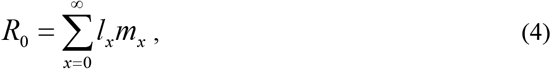

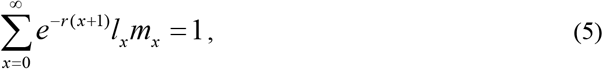

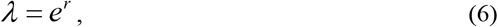

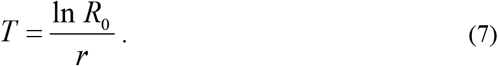

The life expectancy (*e_xj_*), i.e. the time that an individual of age *x* and stage *j* is expected to live, was calculated according to Chi & Su (2006) as

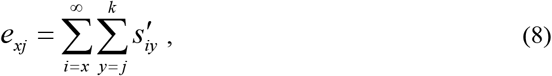

where *s′_iy_* is the probability that an individual of age *x* and stage *j* would survive to age *i* and stage *y*. Fisher (1993) defined the reproductive value (*v_xj_*) as the contribution of individuals of age *x* and stage *j* to the future population. It was calculated as

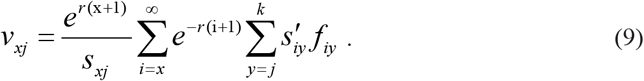

TWOSEX-MSChart software was used to estimate the mean values and standard errors of the population parameters, mean longevity of first instar to fourth instar nymphs and adults, adult and total pre-ovipositional period, mean fecundity per female adult, and the bootstrap test method was used to compare the differences in all the treatments (*B* = 100,000; Chi, 2012). All curve graphs were generated using SigmaPlot 12.0.

## Results

### Dose response bioassay for newly hatched nymphs of *A. glycines*

The LC_50_ concentrations for imidacloprid and thiamethoxam were 4.440 (3.672–5.335) mg a.i./L and 7.049 (5.394–8.998) mg a.i./L respectively, whereas their LC_30_ concentrations were 3.114 (2.425–3.757) mg a.i./L and 4.184 (2.850–5.460) mg a.i./L,respectively (Table 1).

**Table 1.** Toxic effects of imidacloprid and thiamethoxam on newly hatched nymphs of *Aphis glycines*

| Insecticide | Concentration (mg a.i./L) (95 % CL)<sup>-1</sup> | | Slope ± SE | $\chi^2$ (df) |
| --- | --- | --- | --- | --- |
|  | LC<sub>50</sub> | LC<sub>30</sub> |  |  |
| Imidacloprid | 4.440 (3.672–5.335) | 3.114 (2.425–3.757) | 3.402 ± 0.469 | 0.464 (5) |
| Thiamethoxam | 7.049 (5.394–8.998) | 4.184 (2.850–5.460) | 2.314 ± 0.339 | 0.502 (5) |

SE = Standard error

### Life history traits

Imidacloprid and thiamethoxam had effects on the development time, longevity, and fecundity of *A. glycines* (Table 2). Compared with that in the control group, exposure to LC_50_ imidacloprid and thiamethoxam resulted in longer first instar development time, adult pre-ovipositional period (APOP), and total pre-ovipositional period (TPOP; *P* < 0.05), but decreased fecundity (*P* < 0.05) and adult longevity (*P* < 0.05).

**Table 2.**
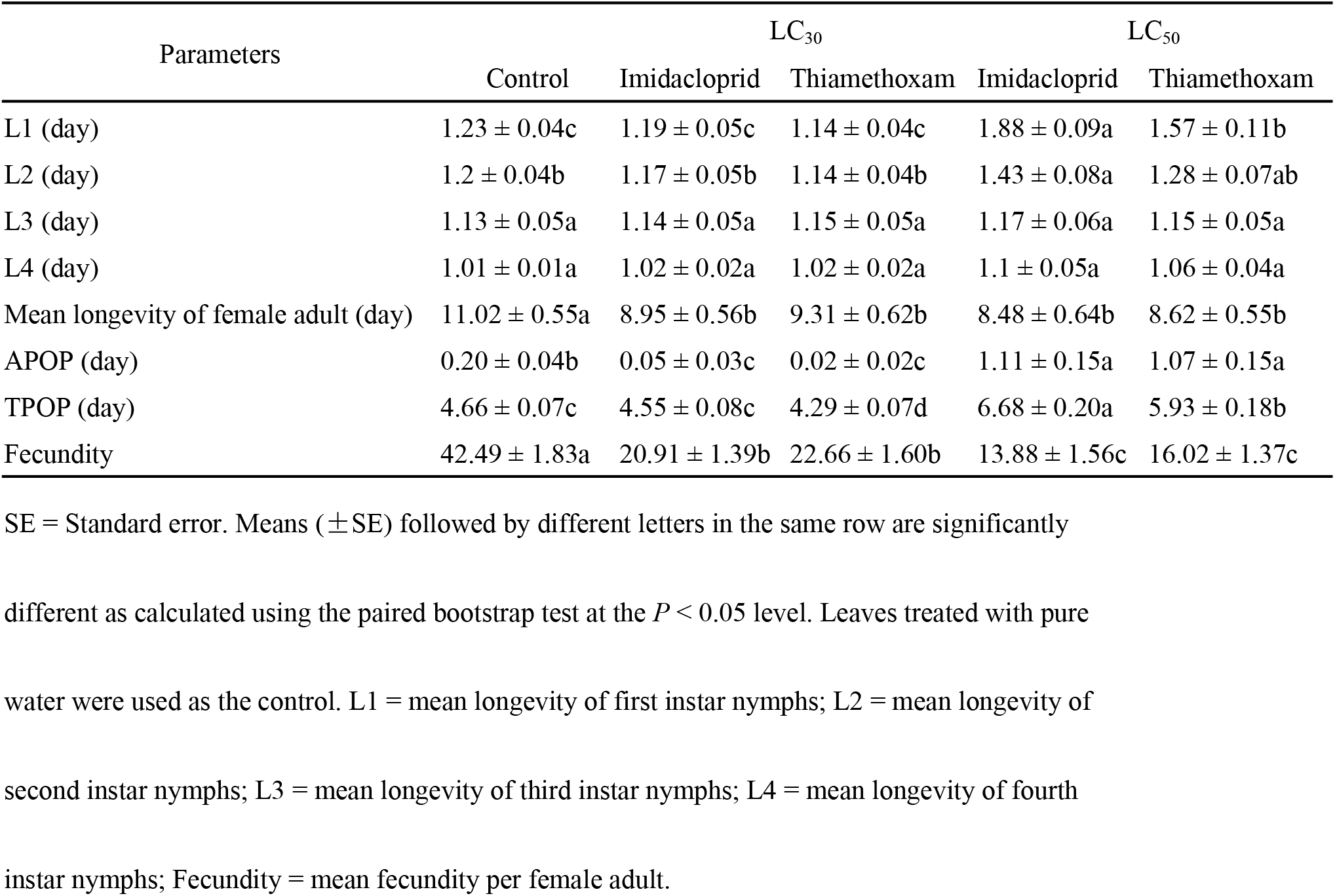
Mean value (± SE) of life history parameters of *Aphis glycines* exposed to imidacloprid and thiamethoxam

Compared with that in the control group (1.20 d), the development time of second instar nymphs increased significantly when exposed to LC_50_ imidacloprid (1.43 d, *P* < 0.05), but no significant change was observed in nymphs exposed to LC_50_ thiamethoxam (1.28 d, *P* > 0.05). The development time of the third and fourth instars did not change on exposure to LC_50_ imidacloprid and thiamethoxam (*P* > 0.05).

Exposure to LC_30_ imidacloprid and thiamethoxam had no significant effect on the development time of the first to fourth instars (*P* > 0.05). However, LC_30_ imidacloprid and thiamethoxam decreased the APOP and reduced the fecundity of *A. glycines* compared with that in the control (*P* < 0.05). The longevity of adults exposed to LC_30_ imidacloprid and thiamethoxam was significantly decreased compared with that of the control group (11.02, 8.95, and 9.31 d in control, imidacloprid, thiamethoxam groups, respectively; *P* < 0.05). The TPOP of *A. glycines* exposed to LC_30_ imidacloprid (4.55 d) showed no significant difference (*P* > 0.05) compared to that in the control group (4.66 d), whereas that of *A. glycines* exposed to LC_30_ thiamethoxam was significantly decreased (4.29 d; *P* < 0.05).

### Life table and fertility parameters

The mean fecundity per female adult, *R*_0_, *r*, and *λ* decreased significantly in the imidacloprid and thiamethoxam treatment groups compared to the control group (*P* < 0.05, Table 3). The *T* in LC_50_ imidacloprid (9.50 d) and thiamethoxam (9.16 d) treatment groups was significantly longer compared with that in the control group (8.20 d; *P* < 0.05). In contrast, the *T* in the LC_30_ thiamethoxam (7.71 d) group was significantly decreased compared with that in the control group (*P* < 0.05), whereas that in the LC_30_ imidacloprid (7.99 d) group showed no significant change (*P* > 0.05, Table 3).

**Table 3.**
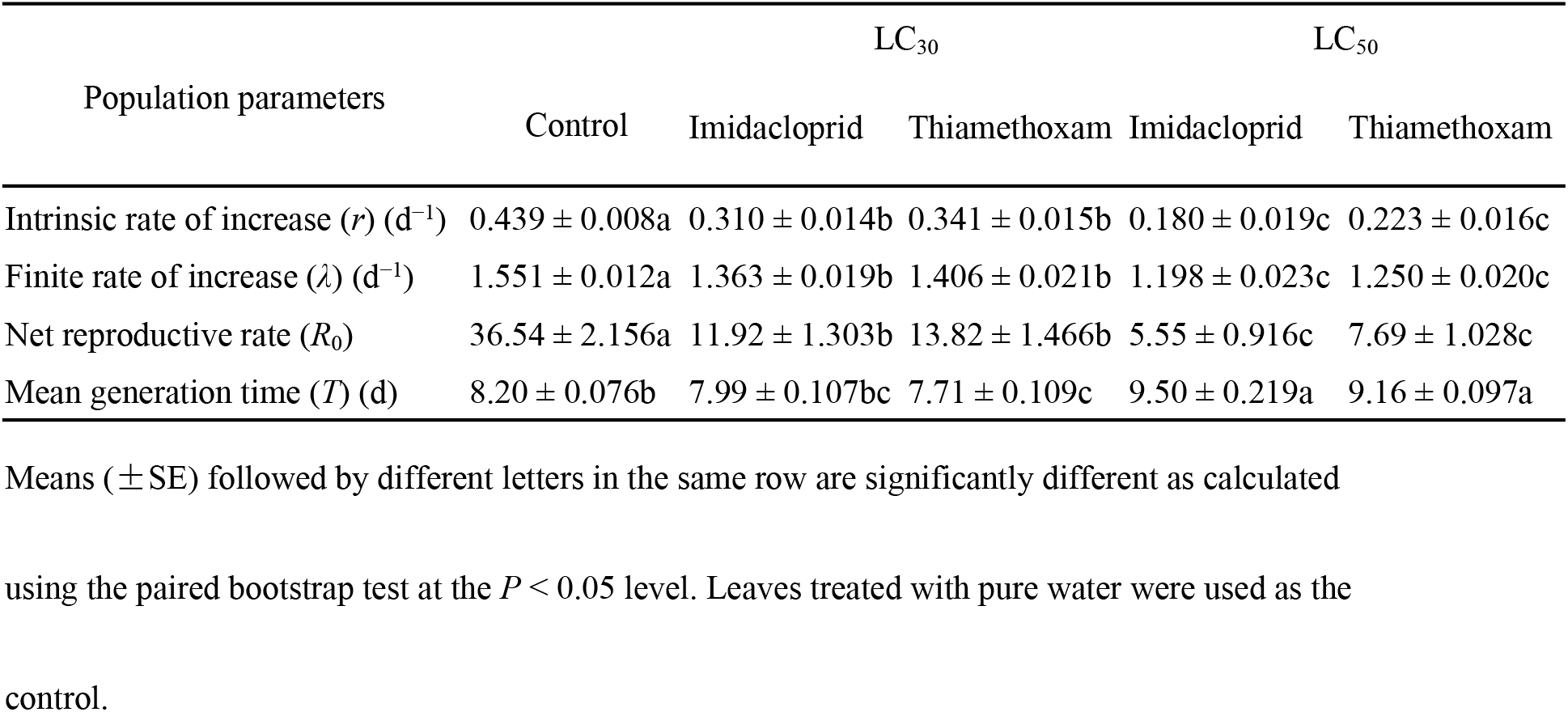
Mean value (± SE) of fertility parameters of *Aphis glycines* exposed to imidacloprid and thiamethoxam

Due to the different development rates between individuals, the age-stage specific survival rates curves show obvious overlaps (Fig 1). The relative number of female adults in the LC_30_ imidacloprid and thiamethoxam treatment groups was higher than that in the respective LC_50_ treatment groups.

Age-specific survival rate is the probability that a newly hatched nymph will reach an age *x*, and the curve of the age-specific survival rate is a simplified form of the curve of the age-stage survival rate, disregarding developmental stages. After treatment with imidacloprid and thiamethoxam, the *l_x_* curve decreased significantly (Fig 2).

The highest peak of *m_x_* in the control group was higher than that in LC_30_ and LC_50_ treatment groups.

The highest peak of *m_x_* in the control group appeared on day 8 (Fig 2), whereas that in the LC_30_ imidacloprid group appeared on day 7, i.e., a day earlier than that in the control group. The highest peak of *m_x_* in the LC_30_ thiamethoxam group appeared on day 6, two days earlier than that in the control group. The highest peak of *m_x_* in the LC_50_ imidacloprid group appeared on day 10, two days later than that in the control group, while the highest peak of *m_x_* in the LC_50_ thiamethoxam group appeared on day 9; a day later than that in the control group (Fig 2).

The values of age-specific maternity (*l_x_m_x_*) were significantly dependent on *l_x_* and *m_x_*, and the maximum *l_x_m_x_* values were 8, 8, 9, 7, and 6 d for the control, LC_50_ imidacloprid, LC_50_ thiamethoxam, LC_30_ imidacloprid, and LC_30_ thiamethoxam treatment groups, respectively.

The female reproductive values in the imidacloprid and thiamethoxam treatment groups decreased compared with those in the control group; however, the female reproductive value in the LC_30_ treatment group was higher than that in the LC_50_ group (Fig 3).

The age-stage life expectancy curve (*e_xj_*) is shown in Fig 4. In the curve, the highest peak values of the first to fourth instar nymphs and female adults were lower in the treatment groups compared with the control group.

## Discussion

The life table parameters used herein reflect the total effect of imidacloprid and thiamethoxam on *A. glycines*. We found that imidacloprid and thiamethoxam at LC_50_ significantly increased the APOP and TPOP and significantly decreased the mean fecundity per female adult compared with that in the control group (*P* < 0.05). In contrast, the APOP in individuals exposed to imidacloprid and thiamethoxam at LC_30_ was shorter than that in the control group (*P* < 0.05, Table 2). In addition, according to the results in the age-stage two-sex life table, the *R*_0_, *λ*, and *r* also decreased significantly (*P* < 0.05, Table 3). Collectively, these results indicate that both imidacloprid and thiamethoxam have inhibitory effects on the reproduction of *A. glycines*.

The *l_x_* curve is a basis for the *s_xj_* curve. In this study, we found that the *l_x_* curves of individuals exposed to imidacloprid and thiamethoxam showed a declining trend (Fig 2). During the first stage, *A. glycines* failed to respond effectively when initially exposed to a high dosage of insecticide; consequently, the *l_x_* decreased sharply, and only some surviving individuals entered the second stage. During the second stage, intoxicated aphids refuse to eat or eat in small amounts and spend energy trying to get out of the toxic arena. This was the stage of confrontation between insecticides stress and *A. glycines*. The different effects of imidacloprid and thiamethoxam at LC_50_ and LC_30_ on the life table may be also related to the regulation strategies of the species, such as self-metabolism and detoxification. In the future research, we will be committed to metabolic detoxification, from the physiological indicators and even molecular level to continue understand the deep impact of imidacloprid and thiamethoxam on population dynamics. During the third stage, the survival rate continued to decline, less sharply than in the first stage, but more sharply than in the second stage. During the third stage, the individual longevity may also be one reason for the decrease of the *l_x_*; this is consistent with the decrease in the age-stage life expectancy curve with increasing age (Fig 4).

In the present study, individuals in the LC_50_ thiamethoxam and LC_50_ imidacloprid treatment groups reached their reproductive peaks 1 and 2 days later compared with those in the control group, respectively. In contrast, individuals in the LC_30_ thiamethoxam and LC_30_ imidacloprid treatment groups reached their reproductive peaks 2 and 1 day earlier than the control group, respectively. Different types and doses of insecticides have different biological and ecological effects on pests. More attention should be paid to the increase of pest reproduction caused by low doses of insecticides (Stark, Tanigoshi, Bounfour, & Antonelli, 1997; James & Price, 2002). For example, low-lethal concentration of spinetoram can decrease the developmental time of *Tetranychus urticae* (Acari: *Tetranychidae*) from egg to adult (Wang, Zhang, Xie, Wu, & Wang, 2016). The rapid increase in the number of *A. glycines* individuals during the productive peak and the fast reproduction of species from generation-to-generation indicate that large outbreaks may occur within a short time. This phenomenon of low-lethal concentration promoting rapid reproduction has brought great pressure on the prevention and control of *A. glycines* in the field.

The actual application doses of neonicotinoids in soybean fields in northeast China was higher than LC_50_, but the complex environmental conditions in the fields and the drift of neonicotinoids themselves would make the doses of some pests exposed to less than LC_50_. The risk of low-lethal doses neonicotinoids stimulate the rapid reproduction of field populations would highly possible happen. We will continue to study whether field populations show similar trends after exposure to low-lethal doses neonicotinoids in the future.

## Supporting information

**S1 Fig.**
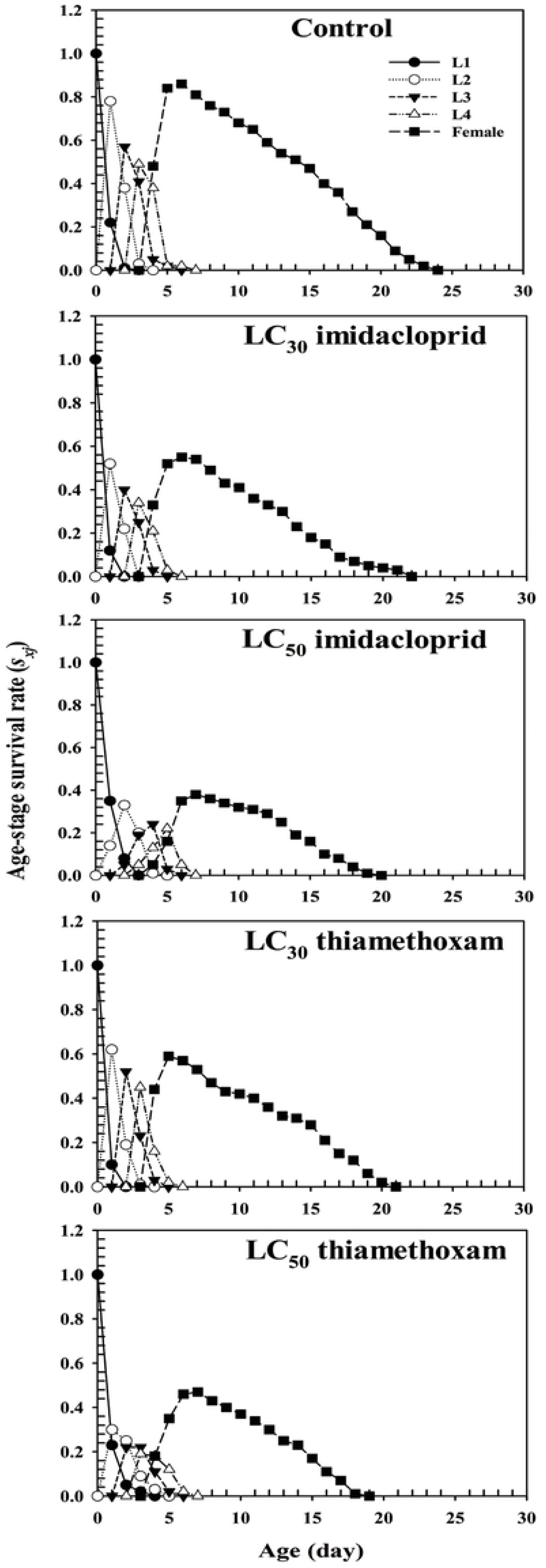
Age-stage specific survival rate (*s_xj_*) of *Aphis glycines*. Age-stage specific survival rate (*s_xj_*) of *Aphis glycines* exposed to the following treatments: control, imidacloprid LC_30_ treatment group, imidacloprid LC_50_ treatment group, thiamethoxam LC_30_ treatment group, and thiamethoxam LC_50_ treatment group. L1 = *s_xj_* of first instar nymphs; L2 = *s_xj_* of second instar nymphs; L3 = *s_xj_* of third instar nymphs; and L4 = *s_xj_* of fourth instar nymphs. (XLS)

**S2 Fig.**
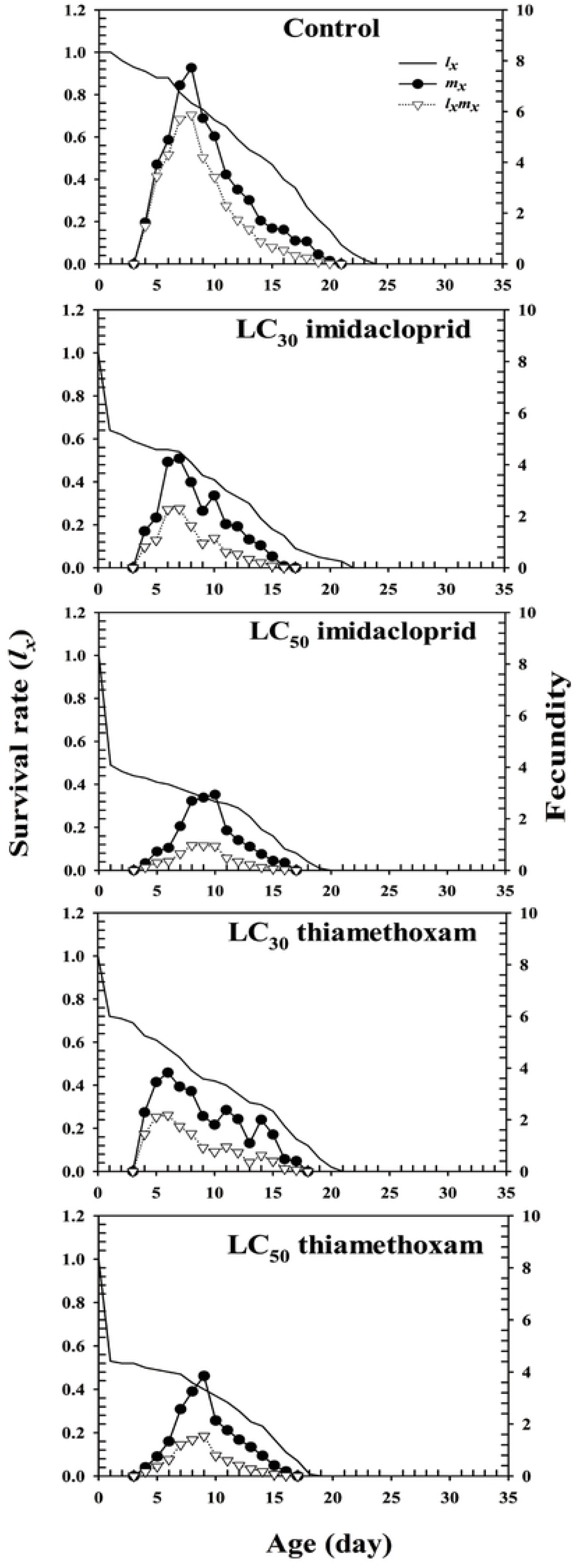
Age-specific survival rate (*l_x_*), age-specific fecundity of the total population (*m_x_*), and age-specific maternity (*l_x_m_x_*) of *Aphis glycines*. Age-specific survival rate (*lx*), age-specific fecundity of the total population (*m_x_*), and age-specific maternity (*l_x_m_x_*) of *Aphis glycines* exposed to the following treatments: control, imidacloprid LC_30_ treatment group, imidacloprid LC_50_ treatment group, thiamethoxam LC_30_ treatment group, and thiamethoxam LC_50_ treatment group. (XLS)

**S3 Fig.**
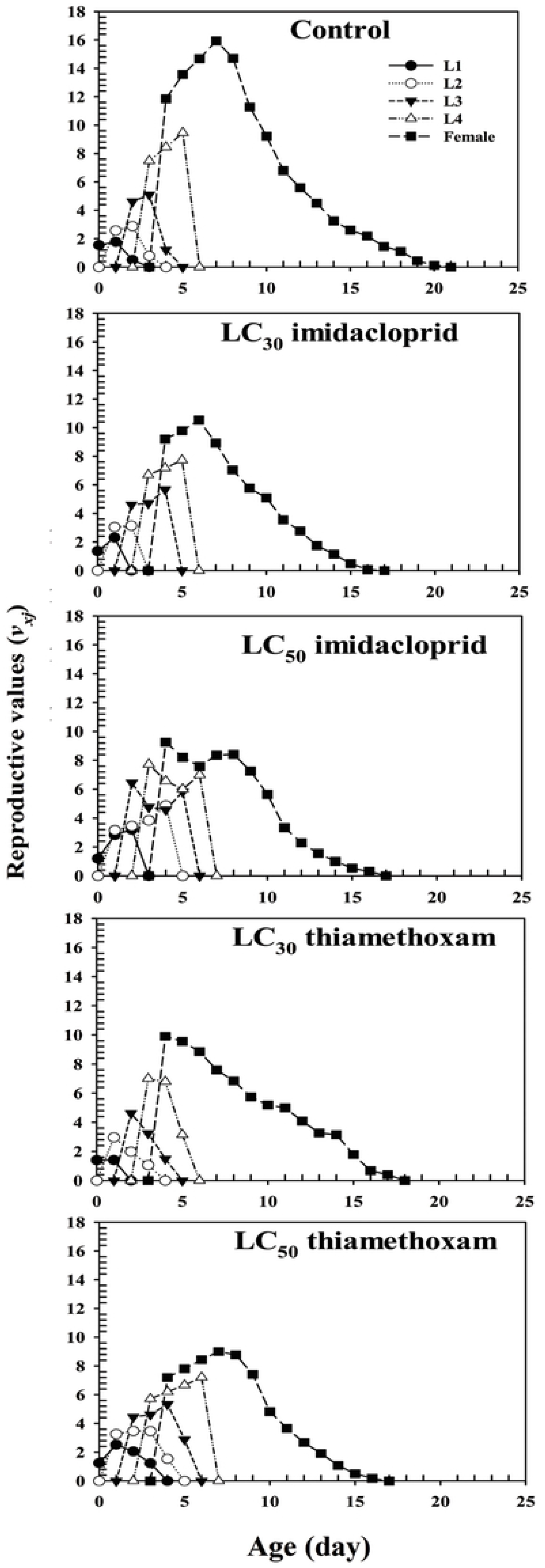
Age-stage specific reproductive values (*v_xj_*) of *Aphis glycines*. Age-stage specific reproductive values (*v_xj_*) of *Aphis glycines* exposed to the following treatments: control, imidacloprid LC_30_ treatment group, imidacloprid LC_50_ treatment group, thiamethoxam LC_30_ treatment group, and thiamethoxam LC_50_ treatment group. L1 = *v_xj_* of first instar nymphs; L2 = *v_xj_* of second instar nymphs; L3 = *v_xj_* of third instar nymphs; and L4 = *v_xj_* of fourth instar nymphs. (XLS)

**S4 Fig.**
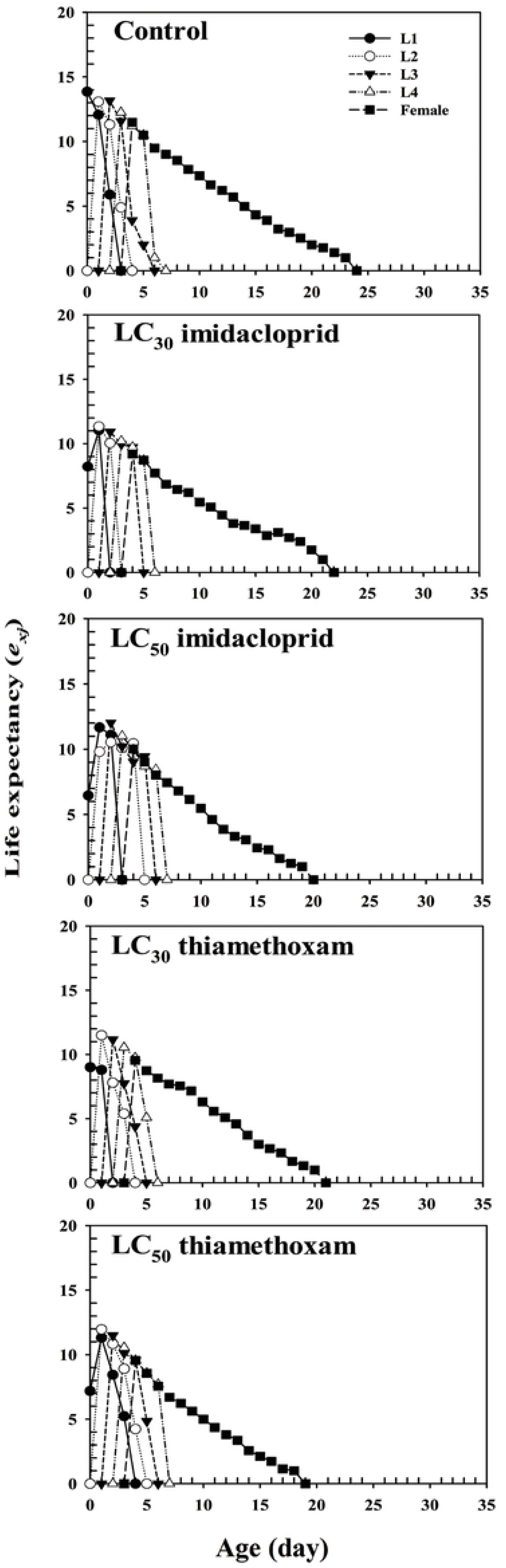
Life expectancy (*e_xj_*) of *Aphis glycines*. Life expectancy (*e_xj_* of *Aphis glycines* exposed to the following treatments: control, imidacloprid LC_30_ treatment group, imidacloprid LC_50_ treatment group, thiamethoxam LC_30_ treatment group, and thiamethoxam LC_50_ treatment group. L1 = *e_xj_* of first instar nymphs; L2 = *e_xj_* of second instar nymphs; L3 = *e_xj_* of third instar nymphs; and L4 = *e_xj_* of fourth instar nymphs. (XLS)

S1 Table. Toxic effects of imidacloprid and thiamethoxam on newly hatched nymphs of *Aphis glycines*. SE = Standard error (XLS)

S2 Table. Mean value (± SE) of life history parameters of *Aphis glycines* exposed to imidacloprid and thiamethoxam. SE = Standard error. Means (± SE) followed by different letters in the same row are significantly different as calculated using the paired bootstrap test at the *P* < 0.05 level. Leaves treated with pure water were used as the control. L1 = mean longevity of first instar nymphs; L2 = mean longevity of second instar nymphs; L3 = mean longevity of third instar nymphs; L4 = mean longevity of fourth instar nymphs; Fecundity = mean fecundity per female adult. (DOC)

S3 Table. Mean value (± SE) of fertility parameters of *Aphis glycines* exposed to imidacloprid and thiamethoxam. Means (± SE) followed by different letters in the same row are significantly different as calculated using the paired bootstrap test at the *P* < 0.05 level. Leaves treated with pure water were used as the control. (DOC)

## Acknowledgments

This work was supported by the Heilongjiang Science Foundation Project (grant number C2018011), Special Fund for Construction of Modern Agricultural Industry Technology System (grant number CARS-04), and Research and Development of Technology and Products on Nature Enemy Insects Prevent and Control (No: 2017YFD0201000).

## Author Contributions

All authors conceived research. AZ conducted experiments. AZ, LZ and TL analysed data and conducted statistical analyses. AZ, LH and ZS wrote the manuscript. LH and KZ secured funding. All authors read and approved the manuscript.

